# Genome comparison reveals inversions and alternative evolutionary history of nutritional endosymbionts in planthoppers (Hemiptera: Fulgoromorpha)

**DOI:** 10.1101/2022.12.07.519479

**Authors:** Junchen Deng, Gordon M. Bennett, Diego C. Franco, Monika Prus-Frankowska, Adam Stroiński, Anna Michalik, Piotr Łukasik

## Abstract

The evolutionary success of sap-feeding hemipteran insects in the suborder Auchenorrhyncha was enabled by nutritional contributions from their heritable endosymbiotic bacteria. However, the symbiont diversity, functions, and evolutionary origins in this large insect group have not been broadly characterized using genomic tools. In particular, the origins and relationships among ancient betaproteobacterial symbionts *Vidania* (in Fulgoromorpha) and *Nasuia/Zinderia* (in Cicadomorpha) are uncertain. Here, we characterized the genomes of *Vidania* and *Sulcia* from three *Pyrops* planthoppers (family Fulgoridae) to understand their metabolic functions and evolutionary histories. Like in previously characterized planthoppers, these symbionts share nutritional responsibilities, with *Vidania* providing seven out of ten essential amino acids. *Sulcia* lineages across the Auchenorrhyncha have a highly conserved genome but with multiple independent rearrangements occurring in an early ancestor of Cicadomorpha or Fulgoromorpha and in a few succeeding lineages. Genomic synteny was also observed within each of the betaproteobacterial symbiont genera *Nasuia*, *Zinderia*, and *Vidania*, but not across them, which challenges the expectation of a shared ancestry for these symbionts. The further comparison of other biological traits strongly suggests an independent origin of *Vidania* early in the planthopper evolution and possibly of *Nasuia* and *Zinderia* in their respective host lineages.

**Originality-Significance Statement:** We sequenced and characterized the genomes of two ancient nutritional symbionts, *Sulcia* and *Vidania*, in three species from the genus *Pyrops* in the species- and symbiont-rich but understudied insect clade, Fulgoromorpha (planthoppers). We describe—for the first time—several independent genome rearrangements in *Sulcia*, which is often cited as a premier example of extreme genome stability spanning hundreds of millions of years. We also show a global lack of synteny across the genomes of the Auchenorrhynchan betaproteobacterial symbionts (*Vidania*, *Nasuia*, and *Zinderia*). This result is unexpected given previous hypotheses of a common origin for these symbionts >250 million years ago alongside *Sulcia*. Taken together, we suggest an independent origin of *Vidania* and possibly of *Nasuia* and *Zinderia* symbiont lineages as well. This hypothesis further links the potential acquisition of novel nutritional endosymbiont lineages with the emergence of auchenorrhyncham superfamilies.

## Introduction

Symbiosis, defined as the long-term interaction between different organisms living closely together (McCutcheon, 2021), plays a critical role in the evolution of life on Earth. The most intimate are endosymbioses where one of the partners lives within the cells of another. The best-known example is the endosymbiosis between an alphaproteobacterium and an archaeon that originated around two billion years ago, where the former partner evolved into what we now know as the mitochondrion within a eukaryotic cell (Imachi *et al*., 2020). Among the different types of eukaryote-bacterial endosymbiosis that evolved subsequently, obligate nutritional endosymbioses have emerged as fundamental drivers of animal biodiversity, particularly in insects (McCutcheon *et al*., 2019). Like mitochondria, obligate endosymbionts undergo strict vertical transmission from mother to offspring and depend on their hosts for many essential cellular functions. Although some of these symbiotic relationships seem to have started relatively recently (Husnik and McCutcheon, 2016; Michalik *et al*., 2021), others have lasted for tens or even hundreds of millions of years (Baumann, 2005; Bennett and Moran, 2015).

A common feature of these ancient relationships is the extremely small genome of the microbial symbiont. The genomes of the beneficial endosymbionts generally undergo rapid and substantial gene loss due to the relaxation of selection on redundant and non-essential genes, elevated mutation rates, population bottlenecks during the vertical transmission, and strong genetic drift that drives the fixation of deleterious mutations even in essential genes (McCutcheon *et al*., 2019). In extreme cases, genome size becomes so small, barely over 100 kilobases, that only functionally essential genes remain, including those involved in transcription, translation, and nutritional provisioning (Bennett and Moran, 2013). As these evolutionary processes proceed, even genes that seem essential in these tiny genomes can be lost over time (Vasquez and Bennett, 2022). Nevertheless, it is typically observed that the genomes of mutualistic endosymbionts establish a high level of synteny fairly early in their evolution (Patiño-Navarrete *et al*., 2013; Williams and Wernegreen, 2015; Chong *et al*., 2019) with some caveats (e.g., Sloan and Moran, 2013). Strict vertical transmission and lack of an environmental phase eliminate the opportunity for recombination between symbiont species and strains (Perreau and Moran, 2021). In addition, the lack of transposable elements and the loss of genes involved in DNA recombination during the extreme size shrinkage also makes structural changes less likely to happen (Silva *et al*., 2003; Chong *et al*., 2019).

Among the best-known nutritional endosymbioses are those in the sap-feeding insect clade Auchenorrhyncha (Hemiptera), which comprises cicadas, spittlebugs, planthoppers, leafhoppers, and treehoppers. To overcome the poor nutritional quality of phloem and xylem plant saps, Auchenorrhyncha species depend on intracellular bacterial symbionts for essential amino acids and vitamins. One of these symbionts, *“Candidatus* Sulcia muelleri” (Bacteroidetes; hereafter: *Sulcia*) established in the common ancestors of Auchenorrhyncha ~300 million years ago (Moran *et al*., 2005; Bennett and Moran, 2013; Johnson *et al*., 2018). Along with *Sulcia*, a diversity of co-resident bacterial symbionts have been described in the four superfamilies, including Betaproteobacteria “*Ca*. Vidania fulgoroideae” (*Vidania*) in planthoppers, “*Ca*. Nasuia deltocephalinicola” (*Nasuia*) in treehoppers and leafhoppers, “*Ca*. Zinderia insecticola” (*Zinderia*) in spittlebugs, and an alphaproteobacterium “*Ca*. Hodgkinia cicadicola” (*Hodgkinia*) in cicadas (McCutcheon and Moran, 2010; Bennett and Moran, 2013). *Sulcia* and these co-resident symbionts together provide a complete set of essential amino acids for the hosts. Among these co-symbionts, the betaproteobacteria *Vidania*, *Zinderia*, and *Nasuia* (referred to as beta-symbionts) share a few genomic characteristics (e.g., extremely reduced genomes) and phylogenomic reconstructions suggested that they might have descended from a single ancestral betaproteobacterial lineage (Bennett and Moran, 2013; Koga *et al*., 2013; Bennett and Mao, 2018), which was replaced by the alphaproteobacterial symbiont *Hodgkinia* in the ancestor of cicadas. However, the genome dissimilarity among beta-symbionts and several known problems with phylogenetic reconstructions, such as the extremely reduced genome and the long-branch attraction, make the hypothesis speculative (discussed in Urban and Cryan, 2012; Bennett and Mao, 2018).

Among all Auchenorrhyncha clades, the least-understood are the planthoppers (Fulgoromorpha) and their endosymbionts. Planthoppers are an ecologically diverse group of >12,000 described species grouped into over 20 families; many of them are economically significant as agricultural pests (Urban and Cryan, 2007; Dietrich, 2009). The early evidence of endosymbionts in planthoppers was revealed by the extensive microscopy work of Müller and Buchner, who described the common presence of two endosymbionts, “a-symbiont” and “x-symbiont”, often accompanied by additional bacteria or fungi (Müller, 1940, 1949; Buchner, 1965). These two common symbionts were identified over half a century later as *Sulcia* and *Vidania*, respectively, with the help of broad sampling and molecular markers (Moran *et al*., 2005; Gonella *et al*., 2011; Urban and Cryan, 2012). However, the genomic study of endosymbionts in Fulgoromorpha was a completely white page until 2018 when the first complete genomes of *Sulcia* and *Vidania* from a Hawaiian planthopper *Oliarus filicicola* (family Cixiidae) were published (Bennett and Mao, 2018). They reported unique features in these planthopper symbionts compared to symbionts in other Auchenorrhyncha superfamilies, including a large chromosomal inversion in *Sulcia* and a dominant nutritional role of *Vidania* over *Sulcia*. A recent study confirmed these patterns in three planthopper species from a divergent family Dictyopharidae (Michalik *et al*., 2021), indicating a likely different evolutionary trajectory of endosymbiosis in planthoppers than in other superfamilies. However, with the symbiont genome-level data available for only four species representing two out of over 20 planthopper families, our comprehension of the evolutionary history in this insect group is far from complete.

To better understand the endosymbiosis in Fulgoromorpha, we characterized *Sulcia* and *Vidania* genomes from three planthopper species in the genus *Pyrops* (family Fulgoridae). This genus includes some of the largest and most spectacular planthopper species, often characterized by elongated head and contrasting color patterns on the wings (Urban and Cryan, 2009). We asked 1) how similar *Sulcia* and *Vidania* are to previously published planthopper symbiont strains, using whole genome assembly from high-throughput sequencing data, and comparative genomics approaches. After discovering a novel rearrangement in *Sulcia* genomes from *Pyrops*, we then asked 2) how common genome rearrangements have been in *Sulcia*, and 3) whether genomes organization and function across *Vidania*, *Nasuia*, and *Zinderia* support hypotheses about their shared origin. We show that *Sulcia* and *Vidania* from *Pyrops* planthoppers are highly similar to previously sequenced strains. *Sulcia* lineages have gone through several large genome rearrangements. In contrast, the unexpected lack of synteny between beta-symbiont genomes strongly argues against their shared ancestry. Although our improved phylogeny still placed *Vidania*, *Nasuia*, and *Zinderia* within a highly-supported monophyletic group, the lingering issue of long-branch attraction in these extremely reduced genomes makes the interpretation from phylogeny inconclusive. Taken together, our data favored multiple origins of beta-symbionts, particularly for *Vidania*.

## Experimental Procedures

### Sample processing and metagenomic sequencing

Individuals of three *Pyrops* species (*Pyrops lathburii* [PYRLAN], *P. clavatus* [PYRCLA], and *P. viridirostris* [PYRVIR]) were sampled from their natural habitat in Vietnam in 2019. Bacteriomes were dissected from a single female for each species. We extracted High-Molecular Weight Genomic DNA (HMW DNA) from the dissected bacteriomes with MagAttract HMW DNA Kit (QIAGEN). Metagenomic libraries of three species were prepared with NEBNext Ultra II DNA Library Prep Kit (BioLabs, New England) with a target insert length of 350 bp and sequenced on Illumina NovaSeq 6000 S4 (2×150bp reads). For long-read sequencing, libraries were prepared with Ligation Sequencing Kit (SQK-LSK 109), and sequenced on MinION Mk1C (Oxford Nanopore Technologies) with R9.4.1 Flow Cell.

### Symbiont diversity in bacteriome samples

Initially, we used phyloFlash v3.3 (Gruber-Vodicka *et al*., 2020) to determine the diversity of symbionts by reconstructing the sequences of small subunit ribosomal RNAs (SSU rRNAs; 16S and 18S rRNAs). We then conducted metagenomic assembly (described in the next section) and used anvi-get-sequences-for-hmm-hits from the anvi’o platform (Eren *et al*., 2015) to identify host and symbiont rRNA gene-containing contigs in the Illumina assembly. rRNA sequences were blasted against the NCBI database to determine their taxonomies. We also used NanoTax.py (https://github.com/diecasfranco/Nanotax) to classify contigs into taxonomic groups determined in the previous step. NanoTax.py performs blast searches using assembled contigs against a customized nucleotide database and a protein database containing sequences from previously assembled genomes of hosts, symbionts, and their free-living relatives. Other information, including the GC content, coverage, and length of each contig, is compiled into output with the assigned taxonomy. Contigs identified as representing symbionts and larger than 1500 base-pairs (bp) were plotted with Processing 3 (http://www.processing.org) in Fig. 1B.

**Figure 1.**
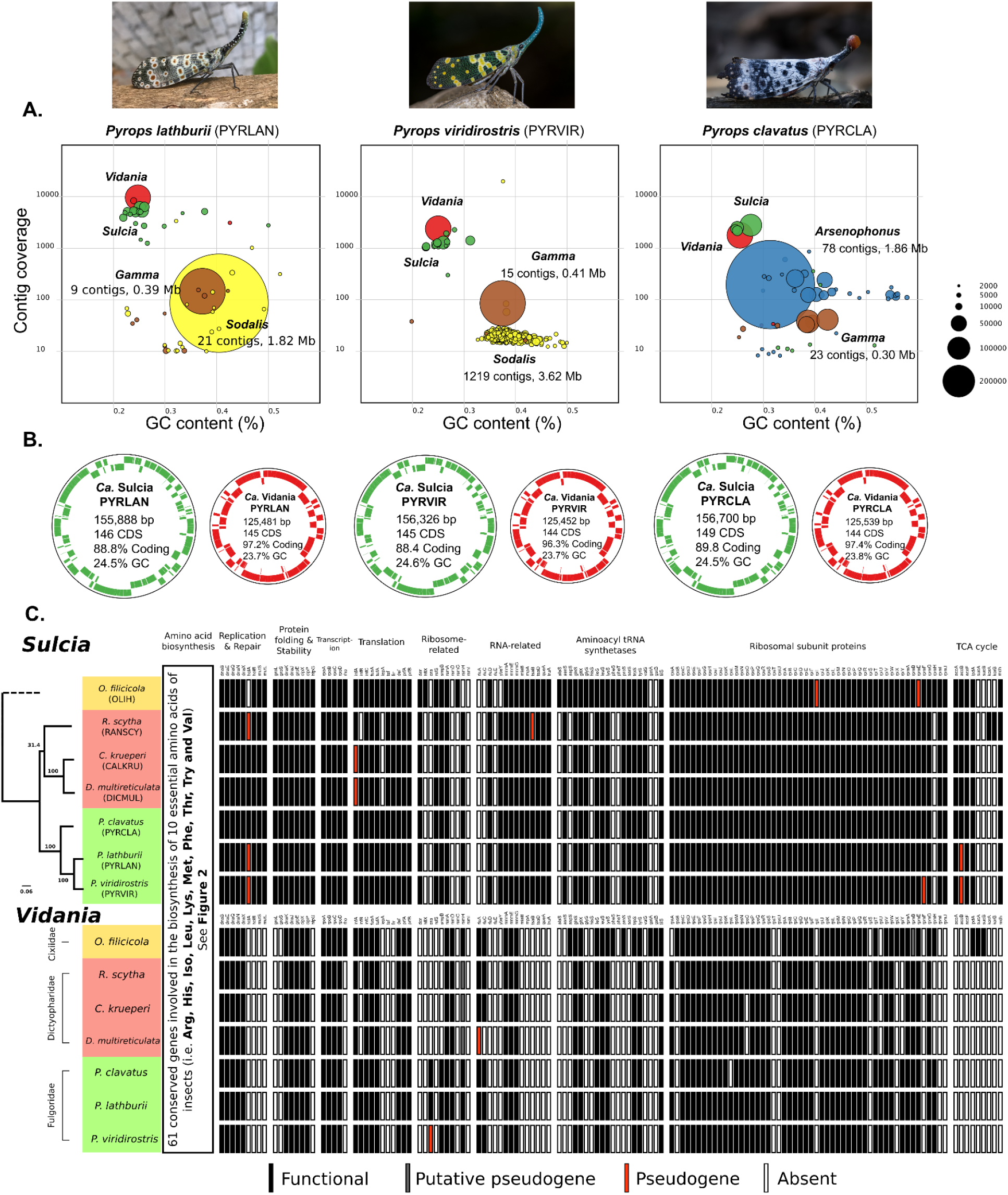
The summary of metagenomic assemblies of three *Pyrops* planthoppers. **A.** The visual representation of endosymbiont diversity based on short-read assembly. Each circle represents a single contig. The circle size is proportional to the length of the contig as it is plotted in GC% and read coverage space. Different colors are applied to different endosymbionts: *Sulcia* (green), *Vidania* (red), *Sodalis* (yellow), *Arsenophonus* (blue) and *Gamma* (brown). Note that *Sulcia* genomes were fragmented in the short-read assemblies but were circular in long-read assemblies, which were then polished by short reads to achieve the final quality. **B.** The visualization of the circular genomes of *Sulcia* and *Vidania*. Genes on forward and reverse strands are shown in colors. Extra information on the genomes, including length (bp), the number of coding DNA sequences (CDS), coding density, and GC content, is shown inside each genome circle. **C.** The comparison of gene set between *Sulcia* and *Vidania* lineages from seven planthopper species representing three families. Each bar represents a single gene with name abbreviation on the top. Genes are classified into functional (black), putative pseudogene (dark gray), pseudogene (red), and absence (white). The maximum likelihood tree of seven host species based on the concatenated ten mitochondrial markers (*nad2*, *cox1*, *cox2*, *atp6*, *cox3*, *nad3*, *nad6*, *cob*, *nad1*, and *rrnL*) is shown on the left. The dotted branch indicates the position of a related species described in the methods section. Insect photographs by Luan Mai Sy, Dr. Vijay Anand Ismavel, and Supratim Deb, originally published on iNaturalist.org.

### Symbiont genome assembly

We conducted metagenomic assemblies as follows: 1) read quality filtering; 2) draft genomes assembly with Nanopore and Illumina reads separately; 3) symbiont genome identification from Nanopore assembly; 4) genome polishing with Nanopore and Illumina reads; and, 5) final genome quality check.

Initially, NanoFilt v2.7.1 (settings: -l 500 --headcrop 10 -q 10) was used to extract high-quality Nanopore reads (De Coster *et al*., 2018). Similarly, trim_galore v0.6.4 (settings: --length 80 -q 30; https://github.com/FelixKrueger/TrimGalore) was used to trim the adapter and control the quality of Illumina paired-end reads. The quality of filtered reads was checked using FastQC v0.11.9 (https://github.com/s-andrews/FastQC). In the second step, Canu v2.1.1 was used to assemble metagenomes from high-quality Nanopore reads (settings: --genome-size 5m and other default settings; Koren *et al*., 2017). Illumina reads were assembled using MEGAHIT v1.1.3 with k-mer size from 99 to 255 (Li *et al*., 2016). In the third step, contigs of *Sulcia*, *Vidania*, and other microbial symbionts were identified using blastn and blastx against custom databases, which included DNA and amino acid sequences of reference genomes of hemipteran insects, mitochondria, microbial symbionts, and their free-living relatives. *Sulcia* and *Vidania* draft genomes, all confirmed circular by canu, were processed further through the polishing step. In the fourth step, high-quality Nanopore reads were trimmed and split using a custom script NanoSplit.py (https://github.com/junchen-deng/NanoSplit), which detects and removes low-quality regions with a sliding-window. All draft genomes were polished by medaka v1.2.1 (https://github.com/nanoporetech/medaka) with the model r941_min_high_g360 for one round and by Pilon v1.23 (Walker *et al*., 2014) using Illumina reads for at least two rounds to fix any mismatches, insertions, and deletions.

In the final step, the polished genomes’ quality was checked by mapping reads to each genome. The original Nanopore reads were filtered by NanoFilt v2.7.1 to recover reads with lengths longer than 1500 bp and an average read quality score >10. Filtered Nanopore reads were then mapped onto polished genomes using minimap2 v2.17 (r941) (Li, 2018). Similarly, the paired-end Illumina reads were mapped onto each genome using bowtie2 v2.4.2 (Langmead and Salzberg, 2012). All mappings were visualized in Tablet (Milne *et al*., 2013) to verify even and consistent coverage (e.g., no breaks).

### Genome annotation

The genomes of *Sulcia* and *Vidania* were annotated with a custom Python script modified from Łukasik et al. (2018). The script first extracted all the Open Reading Frames (ORFs) and their amino acid sequences from a genome. ORFs were searched recursively using HMMER v3.3.1 (Eddy, 2011) against custom databases containing manually curated sets of protein-coding, rRNA, and noncoding RNA (ncRNA) genes from previously characterized *Sulcia* or *Vidania* lineages. rRNA and ncRNA genes were searched with nhmmer (HMMER V3.3.1) (Wheeler and Eddy, 2013), and tRNAs were identified with tRNAscan-SE v2.0.7 (Chan *et al*., 2021). Based on the relative length compared to the reference genes, protein-coding genes were classified as functional (>85%), putative pseudogenes (>60%), or pseudogenes (<60%). Any ORFs over 300 bp but with no significant similarity to any reference genes were blasted against UniProt (The UniProt Consortium *et al*., 2021) and NCBI databases and compared carefully to the top hits. Genes without any annotations were marked as “hypothetical”. All biosynthesis pathways of EAAs were manually constructed with MetaCyc database (Caspi *et al*., 2020).

### Genome comparisons

For the gene set comparison among *Sulcia* and *Vidania* lineages (Fig. 1C), we selected major functional groups that included amino acid biosynthesis, protein folding and stability, replication and repair, transcription, translation, ribosome-related, RNA-related, aminoacyl tRNA synthetases, ribosomal subunit protein, and TCA cycle. Hypothetical genes and genes not involved in any of the above functional groups were not included in the comparison. The genome synteny comparison among *Sulcia* and beta-symbionts (*Vidania*, *Nasuia*, and *Zinderia*) included all published lineages with a recognizable taxonomy available on GenBank (Supporting Information Table S1). The genome synteny comparison among all lineages was illustrated with PROmer v3.07 and MUMmerplot v3.5 (Kurtz *et al*., 2004). Other figures were produced by Processing 3 and were edited in Inkscape.

### Host and Symbiont Phylogenetics

To interpret the results of genome comparison with the host phylogeny, we reconstructed the maximum likelihood phylogeny of host insects based on the concatenated set of ten mitochondrial genes (*nad2*, *cox1*, *cox2*, *atp6*, *cox3*, *nad3*, *nad6*, *cob*, *nad1*, and *rrnL*). The complete mitochondrial genomes of three *Pyrops* planthoppers were assembled and annotated following the methods described in the previous sections. The mitochondrial markers from other host species were extracted from previously published mitochondrial genomes (Supporting Information Table S1). Note that the mitochondrial genomes of *Oliarus filicicola* (OLIH) and *Clastoptera arizonana* (CARI) were unavailable on NCBI, and we replaced them with related species, *Oliarus cf. filicicola* HI01081 and *Philaenus spumarius*, in the analysis. Accordingly, the tree branches of OLIH and CARI were dotted in Fig. 1 and Figs. 3–4. The phylogenetic analyses were conducted in IQ-Tree on XSEDE (Minh *et al*., 2020) implemented in CIPRES v.3.3 (Miller *et al*., 2010). ‘Model Selection’ (Kalyaanamoorthy *et al*., 2017) was selected to search for the best model in CIPRES. The partition type was set to allow the ten partitions (one for each marker) to have different speeds (Chernomor *et al*., 2016). ‘TESTNEWMERGE’ was specified to allow partitions with similar speeds to be analyzed as a single partition. The best fit models were decided by the highest BIC (Bayesian information criterion) scores. Bootstrapping was conducted using ‘SH-aLRT’ bootstrap methods with 1000 replicates. All other setting options were set as default.

**Figure 2.**
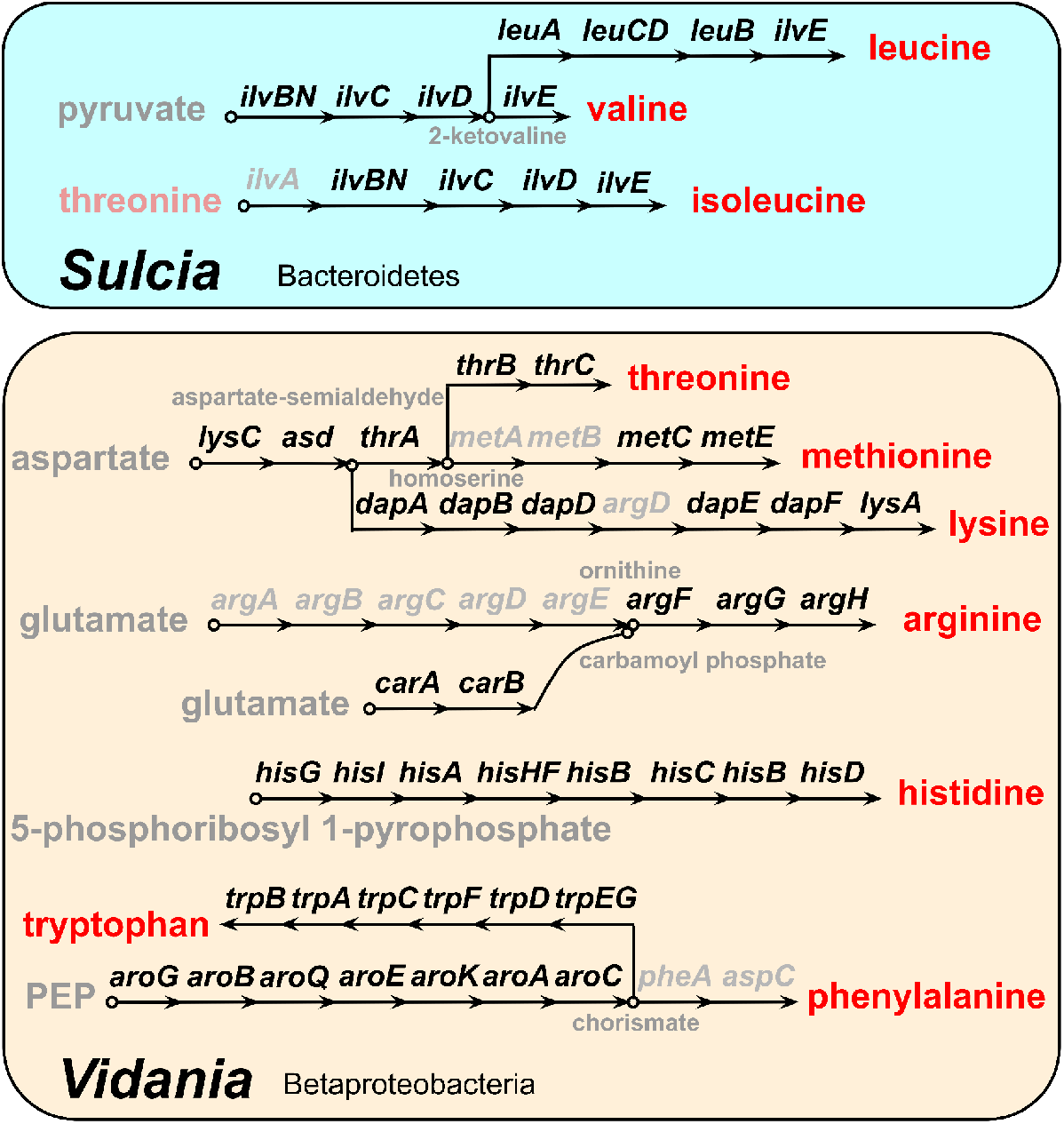
The reconstruction of biosynthesis pathways of ten essential amino acids (EAAs) in *Sulcia* and *Vidania* from three *Pyrops* planthoppers. Each arrow represents a single step in the reaction, with the abbreviated name of the gene shown above. Genes missing from the pathway are shown in gray. EAAs are in red. Chemical compounds involved in some steps are also shown.

**Figure 3.**
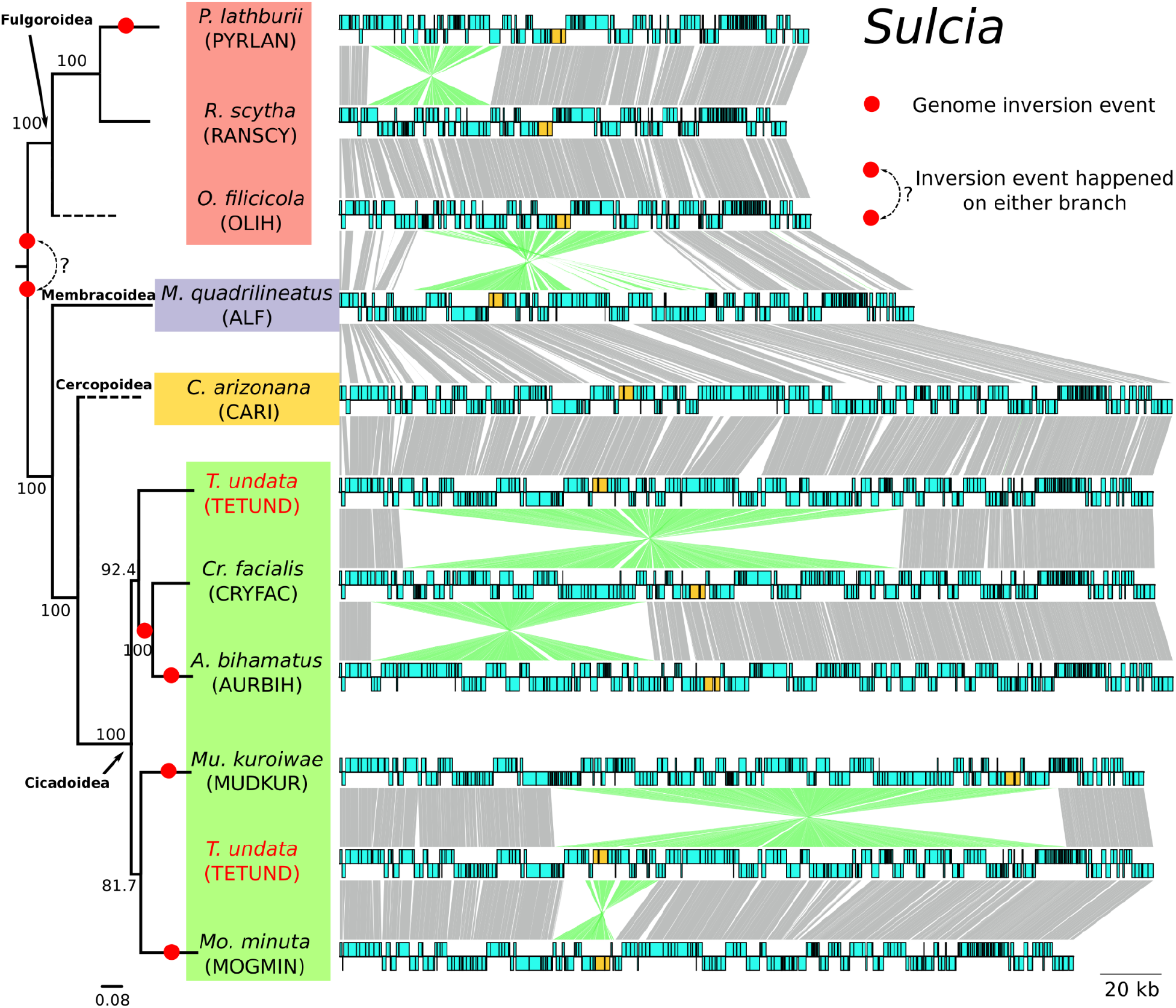
Genome comparison between *Sulcia* lineages. Each circular genome is represented linearly, starting from the same position (*lipB* gene). Genes on the forward and reverse strands are shown on each side as blue boxes. Ribosomal RNAs are colored in orange. Homologous genes are connected by gray lines between genomes. Lines connecting inverted regions are colored in green. The maximum likelihood tree of ten host species based on the concatenated ten mitochondrial markers (*nad2*, *cox1*, *cox2*, *atp6*, *cox3*, *nad3*, *nad6*, *cob*, *nad1*, and *rrnL*) is shown on the left. The dotted branch indicates the position of a related species, as described in the Methods. Note that the lineages shown here represent all inversions we found,. Other genomes with the same organization were omitted from this figure. The comparison of all 41 *Sulcia* lineages can be found in Fig. S1.

**Figure 4.**
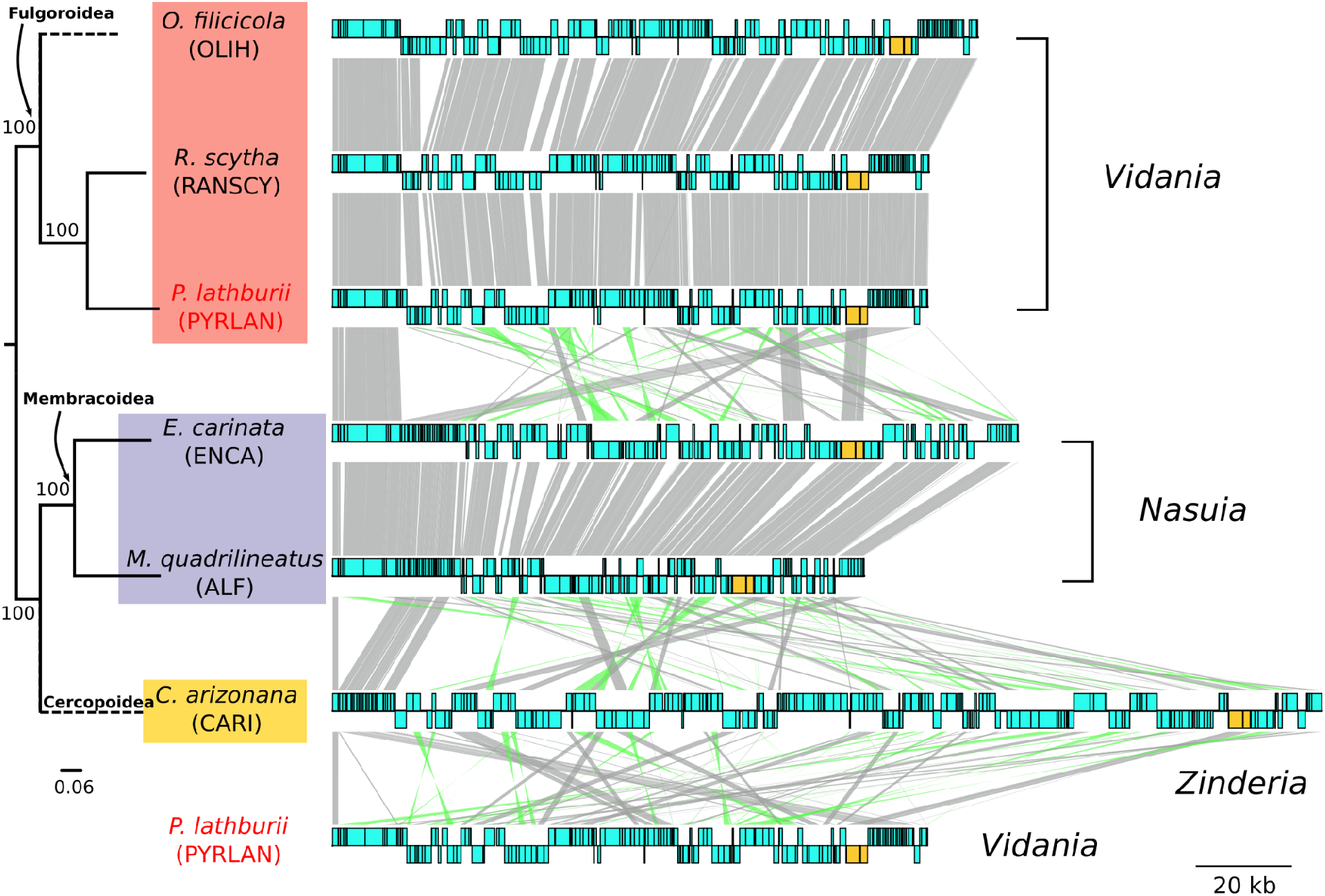
Genome comparison between beta-symbionts (*Vidania*, *Nasuia*, and *Zinderia*). Each circular genome is represented linearly, starting from the same position (*tufA* gene). Genes on the forward and reverse strands are shown on each side as blue boxes. Ribosomal RNAs are colored in orange. Homologous genes are connected by gray lines between genomes. Lines connecting inverted regions are colored in green. The maximum likelihood tree of six host species based on the concatenated ten mitochondrial markers (*nad2*, *cox1*, *cox2*, *atp6*, *cox3*, *nad3*, *nad6*, *cob*, *nad1*, and *rrnL*) is shown on the left. The dotted branch indicates the position of a related species described in the methods section.

To determine the relationships of the sequenced *Betaproteobacterial* strains, as well their placement in the larger betaproteobacterial phylogeny, we reconstructed their relationships using a core-set of protein coding genes. In addition to previously published *Vidania*, *Nasuia*, and *Zinderia* genomes, we extracted gene sequences from unpublished *Vidania* genomes from 23 planthopper species from 12 families (Supporting Information Table S4). Orthologs were determined using HMMER3 searches in Phyloskeleton v.1.1.1 against the 109 bacterial panortholog gene set (settings: *e*-value = 0.01, best-match-only; Darriba *et al*., 2011; Eddy, 2011; Guy, 2017). Genes were translated to amino acid sequences and aligned with Mafft v7 (settings: L-INS-I model; Katoh and Standley, 2013). Models of amino acid substitution were determined with Prottest3 and ambiguously aligned regions trimmed with Trimal v1.4 (Capella-Gutierrez *et al*., 2009; Darriba *et al*., 2011; Supporting Information Data Files S1-2). Trimmed gene alignments were concatenated into a matrix of 289 taxa and 31 genes (7,240 amino acid sites) for phylogenetic analysis (Supporting Information Tables S2-3). RAxML v.8 was used to infer Maximum Likelihood phylogenies from concatenated alignments that were partitioned by gene and run for 500 bootstrap (BS) replicates (Stamatakis, 2014). Two parallel analyses were run for amino acid matrix (-m PTROCATLG) and a Dayhoff6 recoded matrix (-m MULTIGAMMA -K GTR) in an effort to reduce phylogenetic artifacts (Dayhoff *et al*., 1978).

## Results

### Each *Pyrops* planthopper harbors four types of bacteriome-associated endosymbionts

We obtained 11.2, 9.2, and 8.0 Gb of 2×150 bp Illumina data, in addition to 2.5, 1.5, and 1.5 Gb of Nanopore data (N50 = 4.9, 3.2, and 4.7 kb), for three *Pyrops* species (*P. lathburii*, *P. clavatus*, and *P. viridirostris*), respectively. The output of phyloFlash and anvi’o platform confirmed the presence of four distinct endosymbionts in each *Pyrops* species. In each host, we identified the two ancient symbionts, *Sulcia* and *Vidania*, along with two gammaproteobacterial symbionts. These results were confirmed by metagenomic assemblies. The assemblies using short and long reads resulted in complete circular genomes of *Sulcia* and *Vidania* and fragmented genomes of the gammaproteobacterial symbionts.

*Sulcia* and *Vidania* genome sizes range from 155.8-156.7 kb and 125.4-125.6 kb, respectively (Fig. 1B). These sizes are comparable to genomes in family Dictyopharidae (142-148 kb for *Sulcia* and 122-125 kb for *Vidania*) and Cixiidae (157 kb for *Sulcia* and 136 kb for *Vidania*) (Bennett and Mao, 2018; Michalik *et al*., 2021). The Gammaproteobacteria have larger, more complex genomes. All three hosts have a gammaproteobacterial symbiont (denoted as Gamma in Fig. 1A) with relatively small genomes (292-414kb; 9-23 contigs). This symbiont is closely allied to the secondary symbionts of mealybugs and psyllids (Sloan and Moran, 2012; Szabó *et al*., 2017). In addition, *Pyrops lathburii* and *P. viridirostris* have *Sodalis-like* symbionts with genomes of 1.82-3.62 Mb (21-1219 contigs) while *P. clavatus* has an *Arsenophonus* with a genome of 1.86 Mb (78 contigs).

### The conservation of *Sulcia* and *Vidania* genome contents in *Pyrops* planthoppers

The genomes of both *Sulcia* and *Vidania* are highly reduced and gene-dense. Each *Sulcia* genome contains 145-149 predicted protein-coding sequences (CDSs), 29 tRNAs, and a complete ribosomal operon (Fig. 1B, with coding density and GC information). Similarly, each *Vidania* genome contains 144-145 CDSs, 24 to 25 tRNAs, and a complete ribosomal operon (Fig. 1B). Regarding the symbionts’ nutritional roles, *Sulcia* encodes biosynthesis pathways for three EAAs (leucine, valine, and isoleucine) while *Vidania* for seven (methionine, histidine, tryptophan, threonine, lysine, arginine, and phenylalanine) (Fig. 2). Additional gene losses have occurred in several pathways of both *Sulcia* and *Vidania*, including the last step (*pheA* and *aspC*) in phenylalanine biosynthesis, the synthesis of ornithine from glutamate (*argABCDE*) in the arginine pathway, the cysteine pathway (*metAB*) in methionine biosynthesis, and the initial step (*ilvA*) in isoleucine biosynthesis (Fig. 2).

Within the genus *Pyrops*, gene content is highly similar among *Sulcia* and *Vidania*, with only a few differences. *Sulcia* in *P. clavatus* retains the ability to synthesize the subunit delta of the gamma complex of DNA polymerase III (*holA*) involved in DNA replication, as well as the subunit beta of Pyruvate dehydrogenase E1 component (*acoB*) involved in the citric acid cycle (TCA cycle). Both of these genes have been pseudogenized in the other two *Sulcia* lineages (Fig. 1C). In *Vidania*, the gene encoding one ribosomal subunit protein (*rpsL*) was lost from *P. clavatus*.

Gene content between *Pyrops* symbionts and the closely related family Dictyopharidae (RANSCY, CALKRU, and DICMUL; Fig. 1C) is also similar. *Sulcia* in the dictyopharids has lost the translation elongation factor 4 (*lepA*). There was an additional loss of translation initiation factor IF-2 (*infB*) in *Sulcia-RANSCY*. In contrast, *Sulcia-RANSCY* retains the ability to synthesize succinyl-CoA from 2-oxoglutarate through the pathway catalyzed by 2-oxoglutarate oxidoreductase (*korA* and *korB*), which was lost in all other known *Sulcia* lineages from planthoppers. In *Vidania*, translation initiation factor IF-2 (*infB*) was lost in three Dictyopharidae strains.

Finally, compared with symbionts from the more distantly related Cixiidae species, *Oliarus filicicola* (strain OLIH), gene content differences are more pronounced (Fig. 1C). The most notable difference is within the gene set coding for Aminoacyl-tRNA synthetases. *Sulcia* from *Pyrops* and Dictyopharidae have lost five genes for tRNA synthetase (*hisS, pheS, pheT, aspS*, and *alaS*) in contrast to *Sulcia*-OLIH which retains them. Likewise, *Vidania* from *Pyrops* and Dictyopharidae has similarly lost five genes for tRNA synthetases (*aspS, gatA, gatB, pheS*, and *proS*). An additional synthetase loss (*trpS*) occurred in *Vidania-RANSCY*. Furthermore, *Vidania* occurring in *Pyrops* and Dictyopharidae have distinctly lost the complete gene set (*lpd, sucA*, and *sucB*) involved in converting 2-oxoglutarate into succinyl-CoA in the TCA cycle. Two genes involved in the assembly of iron-sulfur clusters (*erpA* and *iscU*, not shown in the figure) responsible for electron transfer were also lost in these two clades.

### Several genome rearrangements have happened in *Sulcia* across Auchenorrhyncha superfamilies

As found previously, *Sulcia-OLIH* (Cixiidae) had a large 78 kb inversion compared to *Sulcia-ALF* from leafhoppers (Bennett and Mao, 2018). *Sulcia* from Dictyopharidae, which is distantly related to cixiids, were collinear to *Sulcia*-OLIH, indicating that this inversion is an ancestral feature of Fulgoromorpha. Here, we observed a novel genome rearrangement (~43 kb including 32 CDSs), in *Sulcia* from *Pyrops* planthoppers compared to cixiid and dictyophorid *Sulcias* (Fig. 3; Supporting Information Figure S1). By further investigating all 41 previously published *Sulcia* genomes across the Auchenorrhyncha, we identified four other instances of genome rearrangements. All of them have occurred in cicadas (Cicadoidea) and were confined to the genera *Auritibicen* (~94 kb, 77 CDSs), *Cryptotympana* (~165 kb, 133 CDSs), *Muda* (~169 kb, 162 CDSs), and *Mogannia* (~20 kb, 16 CDSs) (Fig. 3). In contrast, all *Sulcia* lineages from spittlebugs (Cercopoidea) and leafhoppers and treehoppers (Membracoidea) were collinear and collinear with some cicada *Sulcia* lineages (e.g., *Tettigades undata* TETUND) (Fig. 3; Supporting Information Figure S1). Together, genomes from across the Auchenorrhyncha indicate that an ancestral rearrangement must have occurred in the common ancestor of either Fulgoromorpha or Cicadomorpha shortly after they diverged.

### Betaproteobacterial phylogeny and the lack of synteny between beta-symbiont lineages

Like *Sulcia*, *Vidania* lineages retained a shared gene order (Fig. 4 and Supporting Information Figure S2). This collinearity can also be observed in *Nasuia* (Fig. 4; See also Vasquez and Bennett, 2022). Remarkably, however, the three major beta-symbiont lineages share very little synteny (Fig. 4). The only syntenic regions across all three betaproteobacterial lineages are the more universally conserved ribosomal protein gene clusters (Barloy-Hubler *et al*., 2001).

Results from both amino acid and Dayhoff6-recoded phylogenetic analyses placed the Auchenorrhyncha symbionts into a highly supported monophyletic clade (Fig. 5A; BS = 100; Supporting Information Data Files S3-4). This clade was placed within the Oxalobacteriaceae family (Burkholderiales) as has been found several times previously (BS = 91; e.g., McCutcheon and Moran, 2010; Mao *et al*., 2017). We note that both phylogenetic efforts revealed long-branch attraction between symbionts of scale insects and psyllids insects (e.g., *Tremblaya* and *Proftella*, respectively) and the symbionts of Auchenorrhyncha. These insects are anciently diverged, occurring in the Sternorrhyncha suborder (Hemiptera), and are unlikely to share a common symbiont. Furthermore, the placement of the spittlebug symbiont, *Zinderia* (BS = 100), basal to *Vidania* and *Nasuia* does not match the putative relationships for these major auchenorrhynchan clades (e.g., Johnson *et al*., 2018; Skinner *et al*., 2020). In particular, spittlebugs and leafhoppers are sister groups within the Cicadomorpha; the Cicadomorpha is an early diverging sister clade of the Fulgomorpha planthoppers. This relationship was also recovered in previous phylogenetic efforts, including Bayesian analyses using more sophisticated CAT models (e.g., Bennett and Mao, 2018). These results raise questions about the accuracy of phylogenetic inference within this clade and its potential inability to accurately reconstruct the origins and relationships of betaproteobacterial symbionts. Nevertheless, the relationships of the host species within Fulgomorpha and Cicadomorpha conform closely with our understanding of family, genus, and species (Fig. 5B).

**Figure 5.**
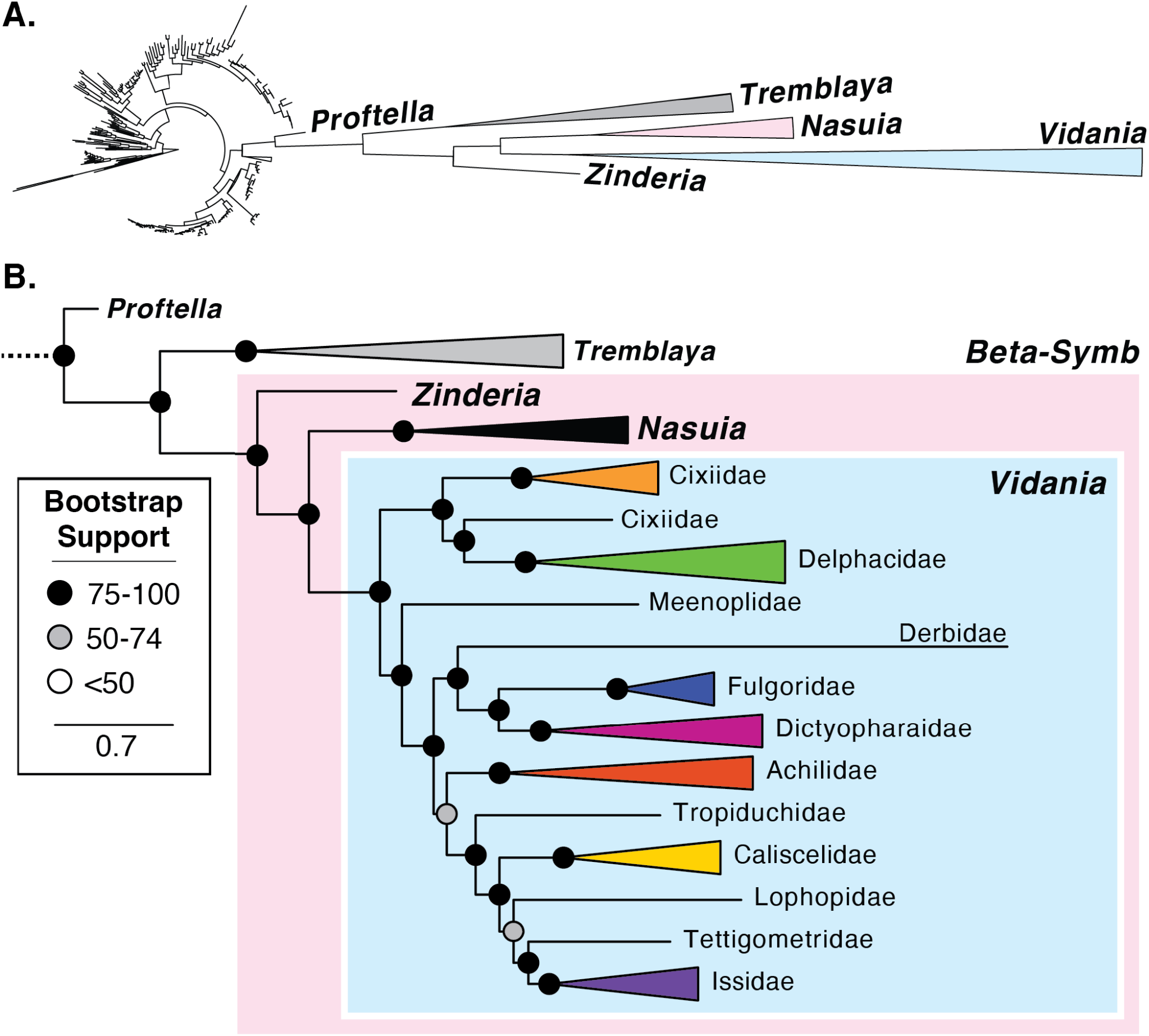
Phylogenetic relationships of the Betaproteobacteria **A.** Maximum likelihood phylogeny reconstructed with 289 taxa and 7240 sites (31 genes). Symbiont clades are labeled. **B.** The zoom-in view of phylogeny showing the relationships among symbionts. *Vidania* clade is collapsed and labeled according to its host family. The full tree can be found in the Supporting Information.

## Discussion

Until recently, Fulgoromorpha has been largely overlooked in the field of endosymbiosis, despite their taxonomic diversity, ecological and economic significance, and diverse endosymbioses that are clearly distinct from those in the sister clade Cicadomorpha (Buchner, 1965; Dietrich, 2009). The newly sequenced genomes of *Sulcia* and *Vidania* from three *Pyrops* planthopper species expand our understanding of the evolutionary history in Fulgoromorpha. The genome of *Sulcia* and *Vidania* of three *Pyrops* planthoppers are highly similar to previously characterized strains, especially from the related Dictyopharidae family. The genome comparison of *Sulcia* lineages revealed that its genome organization has been highly conserved over its ~300 million-year history of co-diversification with the Auchenorrhyncha. Nevertheless, *Sulcia* still experienced genome inversions several times independently: once in the common ancestor of one of the two major auchenorrhynchan clades, and at least five times more recently (Fig. 3). There is no doubt that *Sulcia* is derived from a single ancestral infection that occurred early in the evolution of the Auchenorrhyncha suborder (Moran *et al*., 2005). In contrast, however, the lack of genomic synteny among *Vidania*, *Nasuia*, and *Zinderia* beta-symbionts lineages raises doubts about whether they share a common origin.

### *Sulcia*, *Vidania*, and the additional microbial symbionts in *Pyrops* planthoppers

In *Pyrops* planthoppers, *Sulcia* synthesizes three essential amino acids and *Vidania* provides the remaining seven. The same pattern was observed in four previously investigated planthoppers from families Dictyopharidae and Cixiidae (Bennett and Mao, 2018; Michalik et al., 2021). This pattern is clearly distinct from that in Cicadomorpha, the sister clade of Fulgoromorpha, where *Sulcia* provides more amino acids than its companion (McCutcheon and Moran, 2010; Bennett and Moran, 2013). However, several pathways are incomplete and they lack genes responsible for one or more steps (e.g., arginine biosynthesis in *Vidania*; see fig. 2). These incomplete pathways may be rescued by the host, additional symbionts, moonlighting genes acquiring new functions, or by switching to alternative substrate usage for the nutrient production, as shown in other systems. For example, the loss of the ornithine biosynthesis pathway (*argABCDE*) in arginine biosynthesis, also observed in a few lineages of the nutritional endosymbiont *Buchnera* in aphids (Chong et al., 2019), is compensated by the host using proline rather than glutamate as the substrate for the ornithine synthesis, as suggested by proteomics (Poliakov et al., 2011). The gene losses in other pathways have been fully discussed by Bennett and Mao (2018).

The impressive symbiont diversity in planthoppers has been noted by Müller and Buchner in their classic microscopy work (Müller, 1940, 1949; Buchner, 1965). Recent studies combining modern microscopy techniques and molecular tools suggest that endosymbionts additional to *Sulcia* and *Vidania* may also play significant roles in nutrient provisioning and metabolite exchange. For example, in the family Cixiidae, the gammaproteobacterium *Purcelliella* likely provides B vitamins and metabolic support for *Sulcia* and *Vidania* (Bennett and Mao, 2018). In the family Dictyopharidae, bacteria *Sodalis* and *Arsenophonus*, in the absence of *Purcelliella*, also encode genes involved in B vitamins synthesis (Michalik *et al*., 2021). The highly reduced *Sodalis*-like symbionts and the more recently acquired *Sodalis/Arsenophonus* symbionts in three *Pyrops* species may play similar roles. Their functions will be explored in a separate publication.

### Genome rearrangements in *Sulcia*

The evolution of bacterial endosymbiont genomes is thought to be rapid and chaotic soon after they are acquired by an insect host (McCutcheon *et al*., 2019). Genes critical to the symbiosis are generally under strong selection and are maintained, while others experience relaxed selection and intense genetic drift (Moran, 1996). In endosymbionts that follow strict vertical transmission, these redundant genes are eventually purged from the bacterial genomes (Bobay and Ochman, 2017). The end result is usually a highly stable and compact bacterial genome, as shown in the ancient symbionts of diverse insects (e.g., *Buchnera* in aphids; Chong *et al*., 2019; *Blochmannia* in ants; Williams and Wernegreen, 2015; *Blattabacterium* in cockroaches; Patiño-Navarrete *et al*., 2013; *Wigglesworthia* in tsetse flies; Rio *et al*., 2012). Sulcia is among the most stable symbionts identified so far (McCutcheon *et al*., 2009a; Bennett and Moran, 2013). Indeed, despite substantial differences in gene content and genome size between lineages of Fulgoromorpha and Cicadomorpha, the comparison of *Sulcia* lineages from across the suborder revealed a highly conserved gene order despite its ~300 million years (My) history of co-evolution and restriction to anciently diverged insect lineages.

Despite the overall conserved gene order, several genome rearrangements have occurred in *Sulcia* during its co-diversification with the Auchenorrhyncha. One took place during or soon after the divergence of Fulgoromorpha and Cicadomorpha, and others occurred much later in ancestors of different genera or tribes. These genome inversions were unexpected in a symbiont provided often as an example of genomic stability. However, occasional recombinations and rearrangements have occurred in other ancient endosymbionts of insects. In carpenter ants, the genome comparison between three divergent lineages of the nutritional endosymbiont *Blochmannia*, spanning ~40 My, revealed eight inversions, consisting of two to 34 genes (Williams and Wernegreen, 2015). In a more ancient symbiosis (~140 My) in cockroaches, the comparison between five lineages of the primary endosymbiont *Blattabacterium* revealed three inversions, ranging between 2.9-242 kb (Patiño-Navarrete *et al*., 2013). In aphids, the genomic comparison of 39 strains of the obligate endosymbiont *Buchnera* revealed perfect synteny over 100 My, with the exception of a six-gene inversion shared by a few lineages (Chong *et al*., 2019). These rare genome structural changes reflect somewhat dynamic evolution in those highly stable and tiny genomes.

Genome inversions may be caused by an early wave of mobile elements seen in both recently acquired (e.g., *Serratia;* Manzano-Marín *et al*., 2012; *Sodalis;* Clayton *et al*., 2012) and ancient endosymbionts (e.g., *Portiera* in whiteflies; Sloan and Moran, 2013). Then, these inversions become randomly fixed or purged by genetic drift. Alternatively, genome inversions may also be subject to natural selection. Inversions may bring fitness costs to bacteria by changing gene positions and creating replication-transcription conflicts that alter the gene expression level and mutation rate. These inversions may be negatively selected and weeded out of bacterial populations (Mackiewicz *et al*., 2001; Merrikh *et al*., 2012). On the other hand, inversions may also introduce structural polymorphism when both the ancestral and the inverted genomes exist. Such polymorphism has been observed in at least three endosymbionts, including *Portiera* in whiteflies (Sloan and Moran, 2013), *Tremblaya* in mealybugs (McCutcheon and von Dohlen, 2011), and *Hodgkinia* in cicadas (Łukasik *et al*., 2018; Gordon *et al*., unpublished). Such genomic structural variation could alter the expression pattern of genes involved in host-bacterial symbioses, providing evolutionary flexibility to adapt to environmental challenges (Hughes, 2000; Bennett and Moran, 2015; McCutcheon *et al*., 2019).

### Several lines of evidence support the independent origin of *Vidania*

Like *Sulcia*, the co-resident beta-symbionts in each Auchenorrhyncha superfamily have co-diversified with their hosts for tens and maybe hundreds of millions of years (Moran *et al*., 2005; Bennett and Moran, 2013; Johnson *et al*., 2018). Their genomes have gone through the dynamic phase of pseudogenization and degradation, becoming tiny, compact, and stable. That stability is evident in a comparison between *Nasuia* strains separated by >100 My, or *Vidania* strains separated by ~200 My (Fig. 4; Johnson et al. 2018). It may be expected that beta-symbionts should share similar patterns of synteny if they descended from a single common ancestor as *Sulcia* did, and as has been observed in other ancient insect symbionts (e.g., Buchnera; Chong *et al*., 2019). However, this is not what we found when comparing the genomes of the three genera (Fig. 4). Remarkably, *Vidania, Nasuia*, and *Zinderia* have little to no synteny or chunks of shared gene order. Given the general patterns for symbiont lineages with tiny genomes, this result provides a strong argument that the three beta-symbiont lineages may not be derived from a common ancestral infection. On the contrary, these patterns support the scenario where each of the major auchenorrhychan host lineages that harbor a distinct beta-symbiont lineage (e.g., *Zinderia* in spittlebugs and *Nasuia* in leafhoppers) acquired novel symbionts independently, early in their diversification.

Despite expectations of genomic synteny in ancient symbionts, other evolutionary processes could also result in significant changes in the genome structure of closely related lineages (Sloan and Moran, 2013; Santos-Garcia *et al*., 2020). Specifically, in some clades of whiteflies, their ancient nutritional endosymbiont *Portiera* accumulated repetitive sequences and expanded intergenic regions, causing extensive recombinations and rearrangements in the genome. Interestingly, syntenically expanded genomes were observed in three host lineages separated by at least 7 My, indicating a return to the relatively stable stage after the instability (Santos-Garcia *et al*., 2020). We could envision that a similar genome expansion occurred early in the evolution of beta-symbionts among the major Auchenorrhyncha lineages. However, in the case of *Portiera*, the genome expansion was linked to the loss of the DNA polymerase proofreading subunit (*dnaQ*), which is still retained by all examined lineages of *Vidania*, *Nasuia*, and *Zinderia*. Nevertheless, beta-symbionts lost the DNA mismatch repair system (*mutS* and *mutL*) that has a strong stabilizing effect on the maintenance of genome structure (Nilsson *et al*., 2005), which possibly contributed to the genome instability in the early divergence of beta-symbionts.

We expected that the taxonomically improved phylogeny, including significantly more beta-symbiont genomes, could further resolve the pending issue of beta-symbiont origin. However, the new phylogeny is largely in agreement with previous efforts (Bennett and Moran, 2013; Koga *et al*., 2013; Bennett and Mao, 2018), producing a highly supported clade of the three beta-symbionts. Unfortunately, phylogenetic analyses on those rapidly evolving and extremely reduced genomes all face inevitable phylogenetic errors coming from the taxonomic sampling limitations, extremely elevated rates of molecular evolution, and strong AT nucleotide bias that increase the potential for spurious results like long-branch attraction (Bennett and Mao, 2018). These errors are evident in our analysis as *Tremblaya*, the endosymbiont of distantly related scale insects, was placed within the same clade as Auchenorrhynchan beta-symbionts. In addition, the relationship among *Vidania, Nasuia*, and *Zinderia* in the phylogeny do not agree with the host phylogeny (Cryan and Urban, 2012; Skinner *et al*., 2020). Reconstructing the evolutionary origin of these major beta-symbiont lineages from genomes that have been reduced to <5% of the size of most free-living ancestors is a persistent challenge. It raises the question of whether phylogenetic tools are capable of doing it at all. Hence, we explored whether other evidence (e.g., genomics, metabolisms, bacteriome organization, symbiont cell histology) could support either side of the argument for the origin of beta-symbionts (Table 1).

**Table 1.**
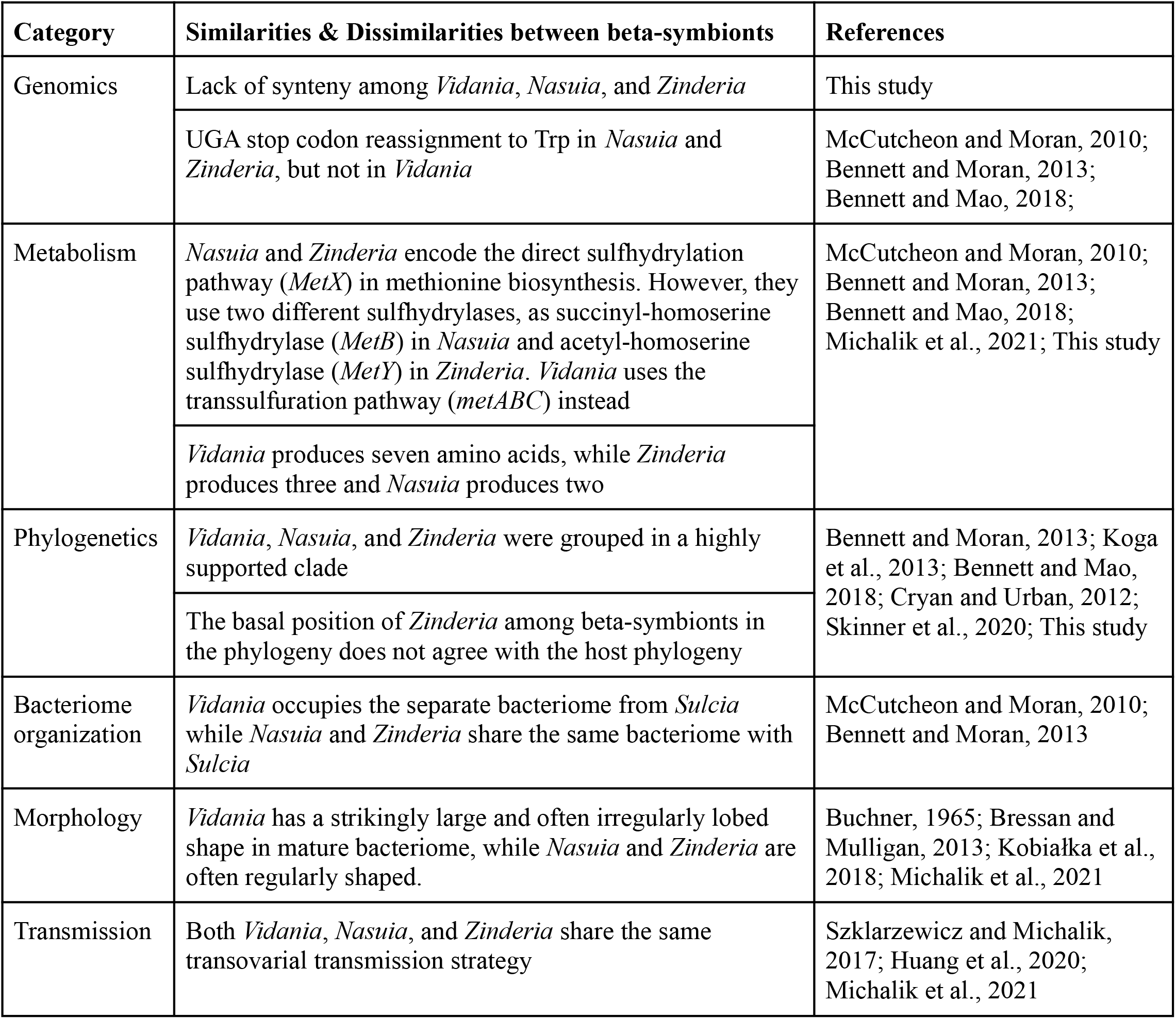
The comparison of biological characteristics between beta-symbionts.

The reassignment of the UGA stop codon to tryptophan in both *Nasuia* and *Zinderia* (McCutcheon and Moran, 2010; Bennett and Moran, 2013), rare in other reduced endosymbiont genomes (Bennett and Moran, 2013), was considered as a strong argument for the close relationship between *Nasuia* and *Zinderia*. They also share the same loss of the peptide chain release factor 2 (*prfB*) that recognizes the UGA stop codon. In contrast, *Vidania* retains the *prfB* gene and does not use the alternative code (Bennett and Mao, 2018). This finding seems to contradict the single ancestry of *Vidania* and *Nasuia/Zinderia*. However, the patterns could be explained by the loss of *prfB* gene in the common ancestor of *Nasuia* and *Zinderia* after its divergence from *Vidania*. Notably, *Hodgkinia*, the independently acquired alphaproteobacterial symbiont in the Cicadoidea superfamily, also lost *prfB* and uses an alternative genetic code (McCutcheon *et al*., 2009a, 2009b). The finding that genetic code reassignment can occur convergently in this group of symbionts of auchenorrhynchan insects weakens argumentation about whether this feature should be given much weight in relationship reconstructions.

More insights into symbiont origins come from the comparisons of how biosynthesis of the ten essential amino acids is partitioned among beta-symbionts and *Sulcia*. In Cicadomorpha, *Sulcia* plays a major role. In spittlebugs and leafhoppers, *Sulcia* provides seven and eight amino acids, respectively, while *Zinderia* and *Nasuia* provide the remaining three and two (McCutcheon and Moran, 2010; Bennett and Moran, 2013). Strikingly, this role is reversed in Fulgoromorpha where *Sulcia* provides only three amino acids while *Vidania* provides seven (Bennett and Mao, 2018; Michalik *et al*., 2021). These patterns are again suggestive of independent origins of Fulgoromorpha and Cicadomorpha symbionts but are not conclusive. The previously characterized multi-symbiont complexes in sap-feeding Hemiptera showed a striking degree of functional complementarity (Husnik and McCutcheon, 2016), indicating that the loss of redundant copies of essential genes generally happens quickly, but stochastic processes could lead to differential gene loss among symbionts in different host lineages originating from a single ancestor. One can imagine that the divergence of Cicadomorpha and Fulgoromorpha occurred soon after the acquisition of *Sulcia* and the beta-symbionts, during which the biosynthetic pathways were lost in a differential manner. On the other hand, however, it would have been surprising if the tryptophan biosynthesis pathway, the one retained by *Zinderia* but not *Nasuia*, was then retained for additional tens of millions of years until the divergence of leafhoppers and spittlebugs.

The details of the methionine biosynthesis pathway also vary among three beta-symbionts. *Nasuia* and *Zinderia* use direct sulfhydrylation pathway (*MetX*) in the production of methionine, but two different sulfhydrylases are used: succinyl-homoserine sulfhydrylase (*MetB*) in *Nasuia* and acetyl-homoserine sulfhydrylase (*MetY*) in *Zinderia* (Bennett and Moran, 2013). In contrast, *Vidania* uses the transsulfuration pathway (*metABC*). The differences in methionine synthesis pathways further point to independent origins for the Fulogmorpha and Cicadomorpha beta-symbionts, further suggesting that *Nasuia* and *Zinderia* are also derived from different original infections. Nevertheless, the different methionine pathways could also be explained by horizontal gene transfer (Gophna *et al*., 2005), or a common ancestor having multiple gene copies, one of which was differentially lost in the major host lineages (Bennett and Moran, 2013).

The internal organization of the host bacteriome organ may also shed light on beta-symbiont origins. Beta-symbionts in two major Auchenorrhyncha clades differ significantly in their bacteriome organization. In Cicadomorpha, Sulcia and beta-symbionts colonize different regions of a single bacteriome (Buchner, 1965; Noda *et al*., 2012; Koga *et al*., 2013; Łukasik *et al*., 2018). This contrasts to Fulgoromorpha, where each endosymbiont is confined to its own, physically distinct bacteriome organ (Buchner, 1965; Bressan and Mulligan, 2013; Michalik *et al*., 2021). The difference in bacteriome organization between the major lineages suggests that symbionts in the two clades have independent origins, infecting different Fulgomorpha and Cicadomorpha host tissues that are later established as bacteriome organs. Nevertheless, the difference in bacteriome organization could also simply reflect the evolution of co-dependency among symbiont species. Symbionts living in adjacent bacteriocytes are more likely to exchange metabolites directly, which has been proposed in Cicadomorpha (McCutcheon and Moran, 2007; Douglas, 2016). This intimacy could also be seen in Fulgoromorpha, as in the family Cixiidae where the third symbiont *Purcelliella*, often sharing the same bacteriome with *Vidania*, may provide metabolites for completing the methionine synthesis pathway (Bressan and Mulligan, 2013; Bennett and Mao, 2018).

Finally, *Vidania* is known for its unusual morphology: strikingly large and often irregularly lobed cells in the mature bacteriome, clearly different from rod-like and regularly shaped cells of *Nasuia* and *Zinderia* (Buchner, 1965; Bressan and Mulligan, 2013; Kobiałka *et al*., 2018; Michalik *et al*., 2021). Interestingly, *Vidania* has a second morphotype, represented by round and regularly shaped cells, in the rectal organ that is unique to female planthoppers and absent in leafhoppers or spittlebugs. This second morphotype, resembling cells undergoing transovarial transmission, was proposed as an infectious form ready to be transmitted to the progeny (Bressan and Mulligan, 2013; Michalik *et al*., 2021). The mechanisms underlying the morphological changes of *Vidania* are still unclear. Nevertheless, morphology is yet another clear difference between Fulgoromorpha and Cicadomorpha symbionts, supporting their independent origins.

## Conclusion

Considering the number of differences between *Vidania* and the two Cicadomorpha beta-symbionts, genomic, metabolic, and morphological evidence suggests that it is highly unlikely that *Vidania* was derived from the same ancestral infection as *Nasuia/Zinderia*. There are more similarities, but also several important differences, between *Nasuia* and *Zinderia*. At the same time, the alphaproteobacterial and clearly independently acquired symbiont of cicadas, *Hodgkinia*, also has extremely reduced genomes, encodes two amino acid biosynthetic pathways, uses an alternative genetic code, inhabits the same bacteriome as *Sulcia*, and shares the same transovarial transmission strategy (McCutcheon *et al*., 2009a, 2009b; Szklarzewicz and Michalik, 2017; Huang *et al*., 2020; Michalik *et al*., 2021). With the exception of the methionine biosynthesis strategy, *Hodgkinia* is about as similar to *Nasuia/Zinderia* as they are to each other, making it clear that this set of similarities can arise through convergent evolution. Both possibilities of independent acquisition of different symbionts by the ancestors of leafhoppers and spittlebugs, and a single infection subsequently exposed to varying evolutionary pressures and constraints that led to differences in genome organization and function, remain viable hypotheses.

There is growing evidence that the tremendous diversity of symbioses of sap-feeding Hemipterans, and many other insects, was shaped by independent, repeated colonization events, and often serial replacements, by different microbes. These microbes include versatile opportunists similar to *Sodalis praecaptivus* (Clayton *et al*., 2012; McCutcheon *et al*., 2019), or specialized pathogens like *Ophiocordyceps* fungi (Matsuura *et al*., 2018). Following colonization, they undergo dynamic genomic reduction in parallel, sometimes in a convergent manner, complementing the function of other symbionts present in the host (Husnik and McCutcheon, 2016). Unfortunately, due to limited sampling and challenges with phylogenetic reconstructions, we are still far from understanding the nature, dynamics, and evolutionary consequences of these replacements, even for relatively recent symbioses. For those that are hundreds of millions of years old, our standard comparative phylogenetic and phylogenomic tools may not be able to accurately and conclusively reconstruct their histories. The combination of a holistic biological approach with more thorough taxonomic sampling across early divergences will give us the best chance of accurately reconstructing the origin and history of these complex symbioses.

## Acknowledgments

We thank all team members of the Symbiosis Evolution Group at Jagiellonian University for helpful discussions. This project was supported by the Polish National Science Center grants 2017/26/D/NZ8/00799 (to A.M.) and 2018/30/E/NZ8/00880 (to P.Ł.) as well as Polish National Agency for Academic Exchange grant PPN/PPO/2018/1/00015 (P.Ł.). G.M.B is funded by the National Science Foundation Division of Biological Infrastructure Award 2214038.

## Author contributions

JD & PŁ conceived the study. AS provided the insect samples. MP prepared sequencing libraries. DCF provided bioinformatic support and additional *Vidania* genomes. GMB provided additional *Nasuia* genomes and conducted phylogenetic analyses. AM contributed morphological observations and interpretations. JD, PŁ, and GMB analyzed the data and wrote the manuscript. All authors reviewed and agreed on the manuscript.

## Data availability

The genomes of *Sulcia* and *Vidania* from three *Pyrops* planthoppers are available under the BioProject PRJNA821037, PRJNA821196, and PRJNA821197 in NCBI databases. The host mitochondrial genomes are available under the accession numbers ON209295-ON209297 in GenBank. Other data underlying phylogenetic analyses (Data Files S1-4) are available in the GitHub repository (https://github.com/junchen-deng/Supplementary-Material_DengEtal_pyrops_2022).

