## Supporting Information Figure for "Genome comparison reveals inversions and alternative evolutionary history of nutritional endosymbionts in planthoppers (Hemiptera: Fulgoromorpha)"

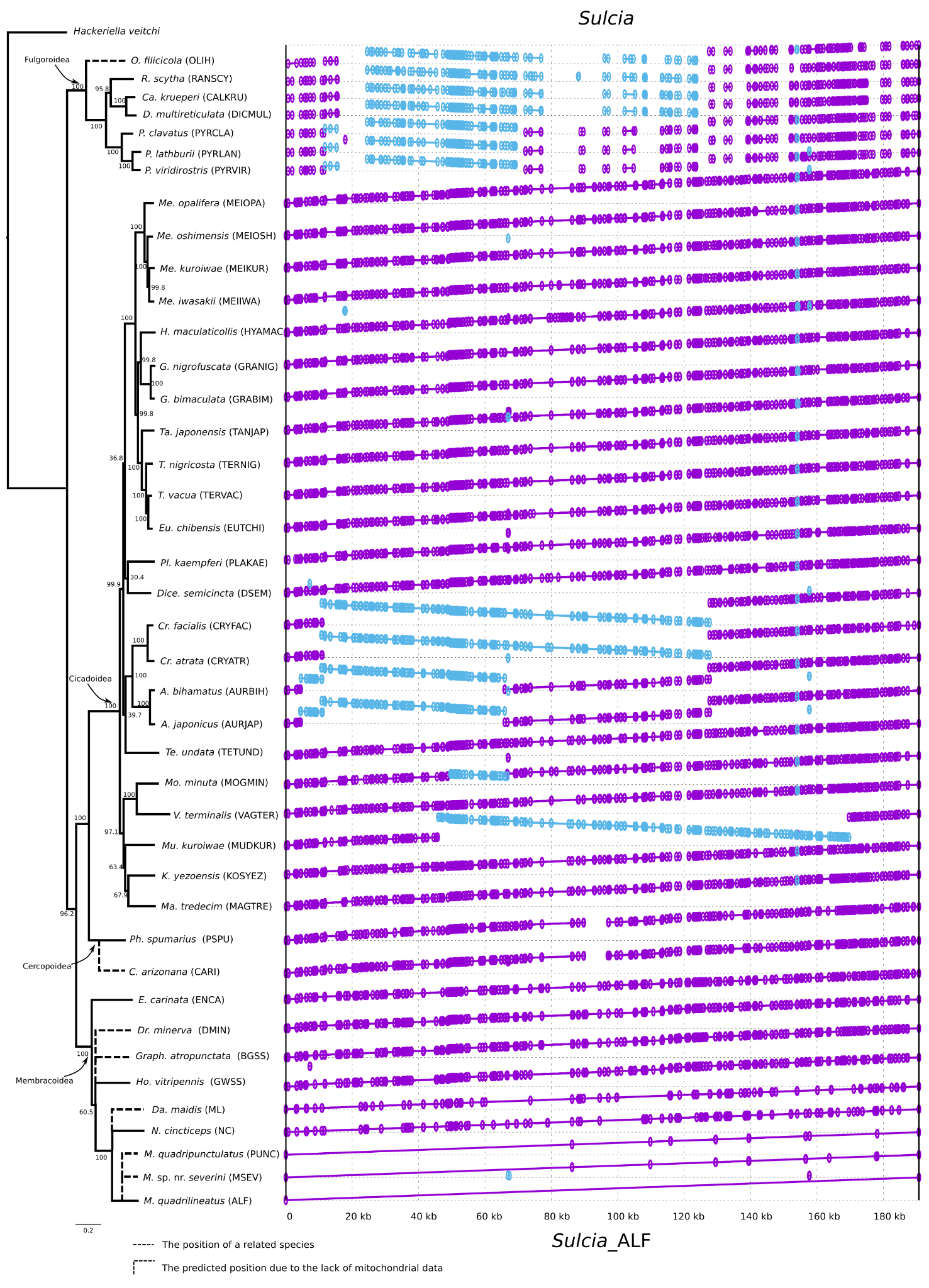

**Figure S1.** The alignment of all *Sulcia* genomes based on the similarity of amino acids. Each genome was aligned with Sulcia from *M. quadrilineatus* (ALF). The purple area indicates the conservation between two genomes. The blue area indicates genome inversions. The gap in the alignment indicates no match or gene loss. The maximum likelihood tree of ten host species based on the concatenated ten mitochondrial markers (*nad2*, *cox1*, *cox2*, *atp6*, *cox3*, *nad3*, *nad6*, *cob*, *nad1*, and *rrnL*) was shown on the left. The Coleorrhyncha species *Hackeriella veitchi* was chosen as the outgroup.

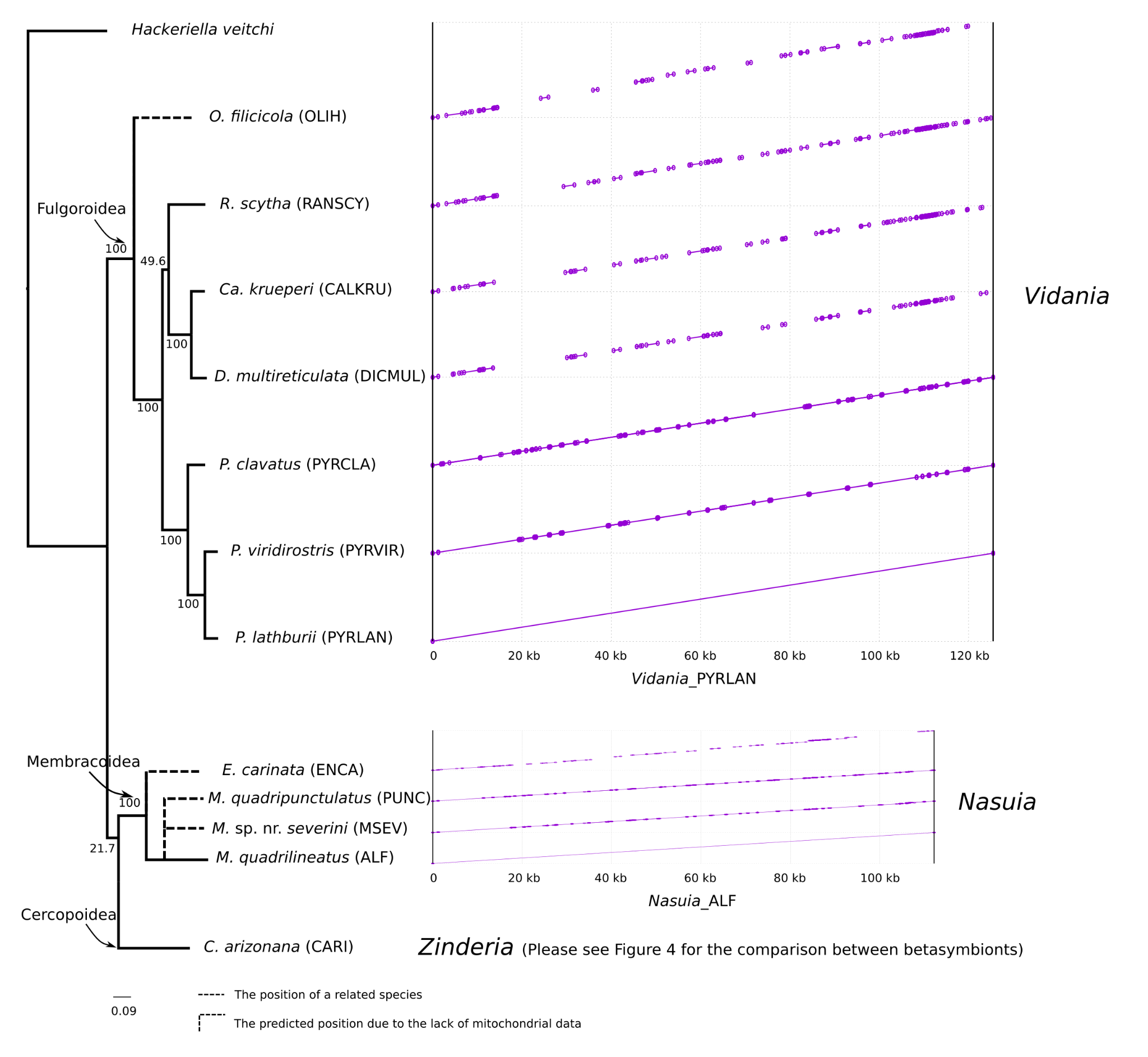

**Figure S2.** The alignment of *Vidania* or *Nasuia* lineages based on the similarity of amino acids. Each *Vidania* was aligned with *Vidania* from *P. lathburii* (PYRLAN). Each *Nasuia* was aligned with *Nasuia* from M. quadrilineatus (ALF). The purple area indicates the conservation between two genomes. The gap in the alignment indicates no match or gene loss. The maximum likelihood tree of ten host species based on the concatenated ten mitochondrial markers (*nad2*, *cox1*, *cox2*, *atp6*, *cox3*, *nad3*, *nad6*, *cob*, *nad1*, and *rrnL*) was shown on the left. The Coleorrhyncha species *Hackeriella veitchi* was chosen as the outgroup.
